# Validating molecular target-enriched fMRI for disentangling drug effects on dopamine

**DOI:** 10.1101/2025.06.27.661944

**Authors:** Ruben van den Bosch, Roshan Cools

**Affiliations:** Radboud University, Donders Centre for Cognitive Neuroimaging, Nijmegen, The Netherlands; Radboud University Medical Center, Department of Psychiatry, Donders Institute for Brain, Cognition and Behaviour, Nijmegen, The Netherlands

**Author notes:** Correspondence: Ruben van den Bosch.

**Keywords:** methylphenidate, dopamine, noradrenaline, fMRI, PET, enriched

## Abstract

The psychostimulant methylphenidate may exert its effects on cognition and associated brain signals via action on the dopamine or noradrenaline transporter (DAT/NET). A recently developed and increasingly popular dual-regression approach (REACT; Dipasquale et al. (2019)) attempts to hone in on the molecular mechanisms underlying (drug-induced) changes in fMRI signal by enriching the analysis with information about the spatial distribution of molecular targets of interest. This method has great potential, but hitherto lacked validation of its molecular specificity and functional relevance. Here we leverage a unique pharmaco-fMRI dataset with established dopamine-dependent methylphenidate effects on neural reward prediction error (RPE) signaling, the canonical functional signature of dopamine. Using REACT we found that methylphenidate significantly modulated both DAT and NET-related functional connectivity networks, but only the effect on the DAT network varied with interindividual differences in striatal dopamine synthesis capacity, as measured with [^18^F]FDOPA PET. Furthermore, methylphenidate affected DAT-related connectivity in the prefrontal cortex in the same location where it affected neural RPE signaling. Together, these findings firmly establish the validity of REACT as a tool for isolating the role of dopamine from that of noradrenaline in methylphenidate’s effects on brain function.

## 1 Introduction

Psychostimulants, like methylphenidate, are widely used for their cognition enhancing effects. Methylphenidate, for example, represents one of the most commonly prescribed treatments for attention-deficit hyperactivity disorder (ADHD), and is also used by healthy people as a party drug or as so-called “smart pills” in the hope of boosting motivation and/or cognitive performance (Maier et al., 2018). Yet, despite more than half a century of methylphenidate prescriptions, we still do not know which of its cognitive effects are mediated by which of its diverse neurochemical mechanisms of action.

What we do know is that methylphenidate acts by inhibiting the reuptake of dopamine and noradrenaline by blocking the dopamine and noradrenaline transporters (DAT and NET, respectively), thereby increasing the bioavailability of dopamine and noradrenaline in the synapse and increasing their signal transmission. The cognitive effects of methylphenidate are thus likely mediated by changes in either dopamine signaling or noradrenaline signaling or both. Unraveling the isolated contribution of dopamine versus noradrenaline to the diverse cognitive effects of methylphenidate is relevant, given that dopamine and noradrenaline are well known to be involved in dissociable aspects of motivation and cognition (del Campo et al., 2011; Varazzani et al., 2015; Westbrook et al., 2020). For example, methylphenidate-related changes in reward learning and motivation for math problem solving might reflect changes in dopamine release in the ventral striatum, as suggested by studies using [^11^C]-raclopride PET imaging (Clatworthy et al., 2009; Volkow et al., 2004). Conversely, methylphenidate’s effects on cognitive variability and sustained attention have been proposed to reflect effects on noradrenaline signaling (Berridge et al., 2012; Milstein et al., 2010).

The most direct way to study the mechanisms of action of psychoactive drugs like methylphenidate in humans is to assess their impact on brain signals with neuroimaging techniques such as functional magnetic resonance imaging (fMRI). However, although such pharmaco-fMRI is well-suited for investigating drug effects on brain-wide activation and network dynamics, it is unable to identify the pharmacological targets that underpin drug-induced changes. The blood-oxygen level dependent (BOLD) signal measured with fMRI has no intrinsic selectivity to any particular neurochemical target. Therefore, network-level brain activity characterized using fMRI remains abstracted from the molecular-level mechanisms through which multi-target drugs in pharmacotherapeutic interventions impart their effects (Lawn, Howard, et al., 2023).

Recently, studies have started to bridge the gap between the micro-scale drug action on molecular targets and the macro-scale drug effects on brain network dynamics, using fMRI analyses informed by the spatial distributions of the molecular targets of interest (Dipasquale et al., 2019; Lawn, Howard, et al., 2023; Salvan et al., 2023). Such integration of modalities has the potential to uncover functional connectivity networks that relate to particular receptors or transporters of interest, e.g. so-called DAT or NET-related functional networks. This promising approach (introduced and dubbed Receptor Enriched Analysis of Connectivity by Targets, or REACT, by Dipasquale et al. (2019)) hones in on the molecular mechanisms underlying changes in BOLD signal while being relatively easy to implement. Any (publicly available) template of molecular target distribution can be used to inform the analysis of (big, publicly available) pharmaco-fMRI data. Accordingly, the method is quickly gaining popularity, with numerous studies linking various neurological and psychiatric disorders, as well as psychoactive drug effects, to specific neurotransmitter-enriched functional connectivity networks (Boucherie et al., 2023; Cercignani et al., 2021; Dipasquale et al., 2020; Lawn et al., 2022; Lawn, Martins, et al., 2023; Luppi et al., 2023; Manca et al., 2025; Puledda et al., 2023; Salvan et al., 2023; Schinz et al., 2023; Wong et al., 2022).

While the approach might have great value, the molecular specificity and functional relevance of the method, for example for isolating the role of dopamine in nonselective drug effects, have not yet been established. Promising results were recently obtained in a study demonstrating that optogenetic activation of serotonin neurons in mice modulates functional connectivity of different large-scale serotonin receptor networks (Salvan et al., 2023), but the evidence for the molecular validity of the large-scale networks derived from the human resting state data in that study remained indirect. It was derived from covariation of neural signal with population-averaged maps of the targets’ spatial distribution, without individual-level molecular imaging data from the same participants. The inference that these large-scale brain networks reflect connectivity with specific molecular targets remains to be substantiated. This is pertinent, because the spatial resolution of MRI is orders of magnitude lower than that of the relevant molecular target distributions.

A separate issue is that the REACT method has so far been used primarily to investigate drug or disease effects on unconstrained BOLD signal measured during resting state. This makes it difficult to estimate the functional relevance of the observed receptor/transporter network changes. This is a serious issue, given that the effects of neuromodulators are known to vary greatly as a function of task demands and neural activity in associated circuits (Cools et al., 2004; Shine et al., 2018; van den Brink et al., 2018). For example, one recent study demonstrated remarkable differences between effects of the serotonergic drug citalopram on receptor-enriched functional connectivity during rest versus task (Boucherie et al., 2023). Diametrically opposite effects on task-related and resting brain states have also been uncovered for the catecholaminergic drug atomoxetine (Shine et al., 2018).

We combined REACT of task-related pharmaco-fMRI with individual-level positron emission tomography (PET) imaging data from the same participants. Our aim was to investigate the molecular mechanisms underpinning the cognitive effects of methylphenidate using a dataset (van den Bosch et al., 2022) that is uniquely qualified to provide functional and molecular validation of the method. This pharmaco-fMRI dataset was obtained from a previous fMRI experiment, in which 94 young health participants underwent fMRI after methylphenidate as well as after placebo, while completing a reversal learning task that has been well established to be sensitive to manipulation of dopamine (Cools et al., 2009; van der Schaaf et al., 2013, 2014). Specifically, in this previous pharmaco-fMRI study (van den Bosch et al., 2022), we used a task that required participants to learn, by trial and error, to predict reward and punishment, given specific stimuli presented to them. Stimulus-outcome contingencies reversed regularly, and reversals were signaled to participants by the presentation of unexpected reward or unexpected punishment. A series of previous studies with this task had already demonstrated that pharmacological changes in dopamine receptor stimulation, e.g. due to withdrawal from dopaminergic medication in Parkinson’s disease (Cools et al., 2006), acute dietary tyrosin depletion (Robinson et al., 2010) or administration of the dopamine receptor antagonist sulpiride in young healthy volunteers (Janssen et al., 2015; van der Schaaf et al., 2014) systematically altered reversal learning from unexpected reward relative to punishment. This dopamine drug-induced asymmetry in reward versus punishment learning concurs with predictions from well-established opponent actor learning models of dopamine’s contribution to learning and decision making that are grounded in the sophisticated push-pull arrangement of direct Go and indirect Nogo pathways of the basal ganglia (Collins & Frank, 2014; Frank, 2005; Jaskir & Frank, 2023). This prior empirical and theoretical work provided the basis for the large pharmaco-fMRI study of methylphenidate’s effects on BOLD signaling in fronto-striatal circuitry. In line with the predictions from this prior work, our previous analysis of this dataset had shown that methylphenidate boosted neural reward- versus punishment-related prediction error (RPE) signals, the canonical functional signature of dopamine, in the anterior and lateral prefrontal cortex (van den Bosch et al., 2022). Importantly, this effect varied with interindividual differences in striatal dopamine synthesis capacity, as measured with [^18^F]FDOPA PET, so that methylphenidate-related increases in neural RPE signaling were greater in participants with higher baseline levels of dopamine synthesis capacity (van den Bosch et al., 2022; cf. van der Schaaf et al., 2013).

Here, we first used REACT to disentangle effects of methylphenidate on the DAT and/or NET-enriched functional connectivity networks, measured during reversal learning task performance. Methylphenidate elicited changes in the DAT-enriched functional connectivity network (as anticipated based on prior REACT work by Dipasquale et al. (2020)), as well as changes in the NET-enriched connectivity network. Critically, methylphenidate’s effects on the DAT network, but not those on the NET network, covaried with (i) interindividual variation in ventral striatal dopamine synthesis capacity as well as with (ii) methylphenidate’s effects on the canonical functional signature of dopamine: neural RPE signaling. These observations substantiate the functional and molecular validity of the REACT tool for disentangling the role of dopamine from that of noradrenaline in the effects of methylphenidate on brain function.

## 2 Methods

### 2.1 Participants

For this study we used data from a previous project (Määttä et al., 2021; van den Bosch et al., 2022) in which 100 healthy volunteers were recruited, 50 women and 50 men (age at inclusion: ranged 18 to 43, mean (SD) = 23.0 (5.0) years). The sample size was determined based on that project’s aim to detect individual differences in dopaminergic drug effects as a function of dopamine synthesis capacity. All participants provided written informed consent and were paid 309 euro after completion of the overarching study. The study was approved by the local ethics committee (“Commissie Mensgebonden Onderzoek”, CMO region Arnhem-Nijmegen, The Netherlands: protocol NL57538.091.16). Prerequisites for participation were an age between 18 and 45 years, Dutch as native language and right-handedness. Before admission to the study, participants were extensively screened for adverse medical and psychiatric conditions, and MRI contraindications. Six participants dropped out of the study, six more were excluded because they failed to meet performance criteria on the reversal learning task, and three datasets were excluded because of poor imaging data quality, resulting in a final sample of N = 85 (van den Bosch et al., 2022).

### 2.2 General procedure

Data were collected as part of a large PET, pharmaco-fMRI study investigating the effects of dopaminergic drugs on brain and cognition, employing a within-subject, placebo-controlled, double-blind cross-over design (Netherlands Trial Register 5959; https://www.onderzoekmetmensen.nl/en/trial/43196). For a detailed description of the testing sessions and tasks and measures collected, see Määttä et al. (2021).

The study consisted of five testing days separated by at least one week. The first was an intake session in which participants were screened for inclusion criteria, an anatomical MRI scan was acquired, and some baseline measures were collected. The second, third and fourth testing days were six-hour-long pharmaco-fMRI sessions in which participants performed a battery of dopamine-related tasks, including the reversal learning task that was performed while being scanned with fMRI. For more information and results regarding the reversal learning task, see van den Bosch et al. (2022). On the fifth day, participants underwent an [^18^F]FDOPA PET scan of the brain to measure their dopamine synthesis capacity.

### 2.3 Image acquisition and preprocessing

The MRI experiment was performed on a 3T Siemens Magnetom Skyra MRI scanner at the Donders Institute, using a 32-channel head coil. Images with blood-oxygen level-dependent (BOLD) contrast were acquired in 3 runs, using a whole-brain T2*-weighted gradient echo multi-echo echo planar imaging (EPI) sequence (38 slices per volume; interleaved slice acquisition; repetition time, 2320 ms; echo times, 9 ms, 19.3 ms, 30 ms, and 40 ms; field of view: 211×211 mm; flip angle 90°; 64×64 matrix; 3.3mm in-plane resolution; 2.5mm slice thickness, 0.4mm slice gap). On the intake session a whole-brain structural MR image was acquired for within-subject registration purposes, using a T1-weighted magnetization prepared, rapid-acquisition gradient echo sequence (192 sagittal slices; repetition time, 2300 ms; echo time, 3.03 ms; field of view: 256×256mm; flip angle, 8°; 256×256 matrix; 1.0mm in-plane resolution; 1.0mm slice thickness).

The brain PET data were acquired on a PET/CT scanner (Siemens Biograph mCT; Siemens Medical Systems, Erlangen, Germany) at the Department of Medical Imaging of the Radboud University Medical Center. We used the well-validated radio-tracer [^18^F]FDOPA, which was synthesized at Radboud Translational Medicine BV (RTM BV) in Nijmegen. The procedure started with a low-dose CT scan to use for attenuation correction of the PET images. Then, the [^18^F]FDOPA tracer was administered (approximately 185 MBq) via a bolus injection in the antecubital vein and the PET scan was started. Dynamic PET data (4×4×3 mm voxel size; 5 mm slice thickness; 200×200×75 matrix) were acquired over 89 minutes and divided into 24 frames (4×1, 3×2, 3×3, 14×5 min). Data were reconstructed with weighted attenuation correction and time-of-flight recovery, scatter corrected, and smoothed with a 3 mm FWHM kernel.

All MRI data were preprocessed using fMRIPrep (1.2.6-1; RRID:SCR_016216; Esteban et al. (2019); Esteban et al. (2020)). Before preprocessing the functional data with fMRIPrep, we combined the multi-echo data into a single time-series per fMRI run with the multi-echo toolbox (https://github.com/Donders-Institute/multiecho; commit Nr.: 9356bc51ef) using the TE algorithm, in which the different echoes are weighed by their echo time. The multi-echo-combined functional scans were then realigned, coregistered to the participant’s T1-weighted anatomical scan, and spatially normalized to MNI152 space with fMRIPrep. More details on the fMRI preprocessing with fMRIPrep can be found in the Supplementary Methods of van den Bosch et al. (2022). We used Statistical Parametric Mapping 12 (SPM12; https://www.fil.ion.ucl.ac.uk/spm/software/spm12/) running in MATLAB R2019b (Mathworks Inc.; https://nl.mathworks.com/products/matlab.html) to spatially smooth the final preprocessed BOLD time-series with a 6 mm FWHM kernel.

The PET data were preprocessed and analyzed using SPM12. All frames were realigned to the mean image to correct for head motion between scans. The realigned frames were then co-registered to the structural MRI scan, using the mean PET image of the first 11 frames (corresponding to the first 24 minutes), which has a better range in image contrast outside the striatum than a mean image over the whole scan time. Presynaptic dopamine synthesis capacity was quantified as the tracer influx rate k_i_^cer^ (min^-1^) per voxel with graphical analysis for irreversible tracer binding using Gjedde-Patlak modeling (Patlak et al., 1983; Patlak & Blasberg, 1985). The analysis was performed on the images corresponding to 24-89 minutes, which is the period after the irreversible compartments had reached equilibrium and the input function to the striatum had become linear. The k_i_^cer^ values represent the rate of tracer accumulation relative to the reference region of cerebellar grey matter, where the density of dopamine receptors and metabolites is extremely low compared with the striatum (Farde et al., 1986; Hall et al., 1999). The cerebellar grey matter mask was obtained using FreeSurfer segmentation of each individual’s anatomical MRI scan, as implemented in fMRIPREP. Resulting k_i_^cer^ maps were spatially normalized to MNI space, smoothed with an 8mm FWHM kernel and brain extracted.

After preprocessing we extracted the mean k_i_^cer^ values from the ventral striatum in native subject space. The striatal striatum mask was obtained from an independent, functional connectivity-based parcellation of the striatum conducted in a previous study (Piray et al., 2017). We focused our analysis on dopamine synthesis capacity in the ventral striatum, given the extensive prior literature demonstrating that methylphenidate acts on dopamine transmission particularly in the nucleus accumbens (e.g. Martinez et al., 2020; Volkow et al., 2002; Volkow et al., 2012) and the fact that methylphenidate’s effect on reward prediction error signaling in the prefrontal cortex varied with dopamine synthesis capacity in the ventral part of the striatum (van den Bosch et al., 2022).

### 2.4 Population-based molecular templates

We used publicly available molecular templates to inform our fMRI analyses with information about the spatial distribution of the density of the transporters of interest. The template for DAT was the averaged DAT map of the healthy controls (N=174) in the Parkinson’s Progression Marker Initiative (https://www.ppmi-info.org/), which used single photon emission tomography (SPECT) with the tracer [^123^I]-FP-CIT. It was obtained from Hansen et al. (2022) (https://github.com/justinehansen/hansen_receptors-1). The template for NET (N=77) was created from PET data using the tracer [^11^C]MRB and was also obtained from Hansen et al. (2022). For our aim to substantiate target-specificity of our findings, we planned to repeat our analyses with a template of the serotonin transporter (SERT). The SERT template (N=100) was created with PET using the transporter [^11^C]DASB and was obtained from Beliveau et al. (2017) (https://xtra.nru.dk/FS5ht-atlas/).

### 2.5 Analysis of molecular target-enriched functional connectivity

For the analysis of the molecular target-related functional connectivity networks we used a dual-regression approach following the method as used in REACT (receptor-enriched analysis of function connectivity by targets; Dipasquale et al. (2019)), implemented in Python and SPM12. First, we resampled the fMRI scans and molecular template atlases to MNI152NLin2009cAsym template space with 2mm isotropic resolution. The reference regions that were used in the creation of the molecular templates (occipital cortex for DAT/NET and cerebellar grey matter for SERT) were masked out of the atlas images, and the atlases were normalized to rescale the images values between 0 and 1. Then, an analysis mask was created from the intersection of the voxels included in the molecular template atlases, a grey matter mask, and those voxels for which fMRI data was available for all participants.

Spatial regressions were performed with one molecular template map and each volume of a participant’s fMRI time series (limited to the voxels in the analysis mask), repeated for each molecular template and for every participant separately. These regressions quantified the correspondence between molecular target density and BOLD signal at each timepoint, resulting in a vector over time with higher values for volumes in which BOLD signal was high in brain regions with high target density. These vectors quantifying the similarity between the spatial map of BOLD signal and the spatial map of the target of interest (i.e. DAT, NET, or SERT) were normalized and added to the general linear models of the regular first-level fMRI analysis for the second regression step. The temporal regression of the spatial similarity regressors in the first-level models estimated the subject-specific spatial maps of target density-enriched functional connectivity, i.e. the DAT/NET-related functional connectivity networks.

Except for the addition of the spatial similarity regressors and the interaction between them and the reversal learning task event regressors (unexpected/expected reward/punishment outcomes), the fMRI analysis was kept exactly the same as previously reported (van den Bosch et al., 2022). By keeping the regressors for task events and their interactions in the model, variance associated with task events was not assigned to the spatial similarity regressors, meaning that the resulting functional connectivity networks were not driven by these task events. This makes the analysis more comparable to previous analyses with resting-state fMRI data (such as Dipasquale et al., 2020).

### 2.6 Statistical analysis

For each participant we contrasted the target-enriched functional connectivity images under methylphenidate with those under placebo. These contrast images were subsequently analyzed with one-sample t-tests to test for methylphenidate effects on target-related functional connectivity at the group-level. We then assessed whether methylphenidate effects on target-enriched functional connectivity varied with interindividual differences in dopamine synthesis capacity by including the [^18^F]FDOPA k_i_^cer^ values of dopamine synthesis rates in the ventral striatum as a covariate. Statistical maps resulting from the analyses were visually assessed using an uncorrected initial cluster-forming threshold of p<0.001, and statistically significant clusters are reported based on cluster-level family-wise error (FWE) correction. Supplementary Table 1 lists the information for all clusters with effects of methylphenidate at p<0.001 uncorrected and/or p<0.05 with whole-brain, cluster-level FWE correction.

## 3 Results

To briefly recap our approach, we assessed methylphenidate’s effects on DAT/NET-enriched functional connectivity networks, by informing our pharmaco-fMRI analyses with publicly available SPECT/PET data on the spatial distribution of these transporters (Figure 1a). Using spatial regression of these molecular templates with the fMRI timeseries we created regressors that quantified the spatial similarity between the pattern of BOLD signal and the DAT/NET density map at each time point. These spatial similarity regressors were then added to the subject-level fMRI models to identify DAT/NET-enriched functional connectivity networks.

**Figure 1.**
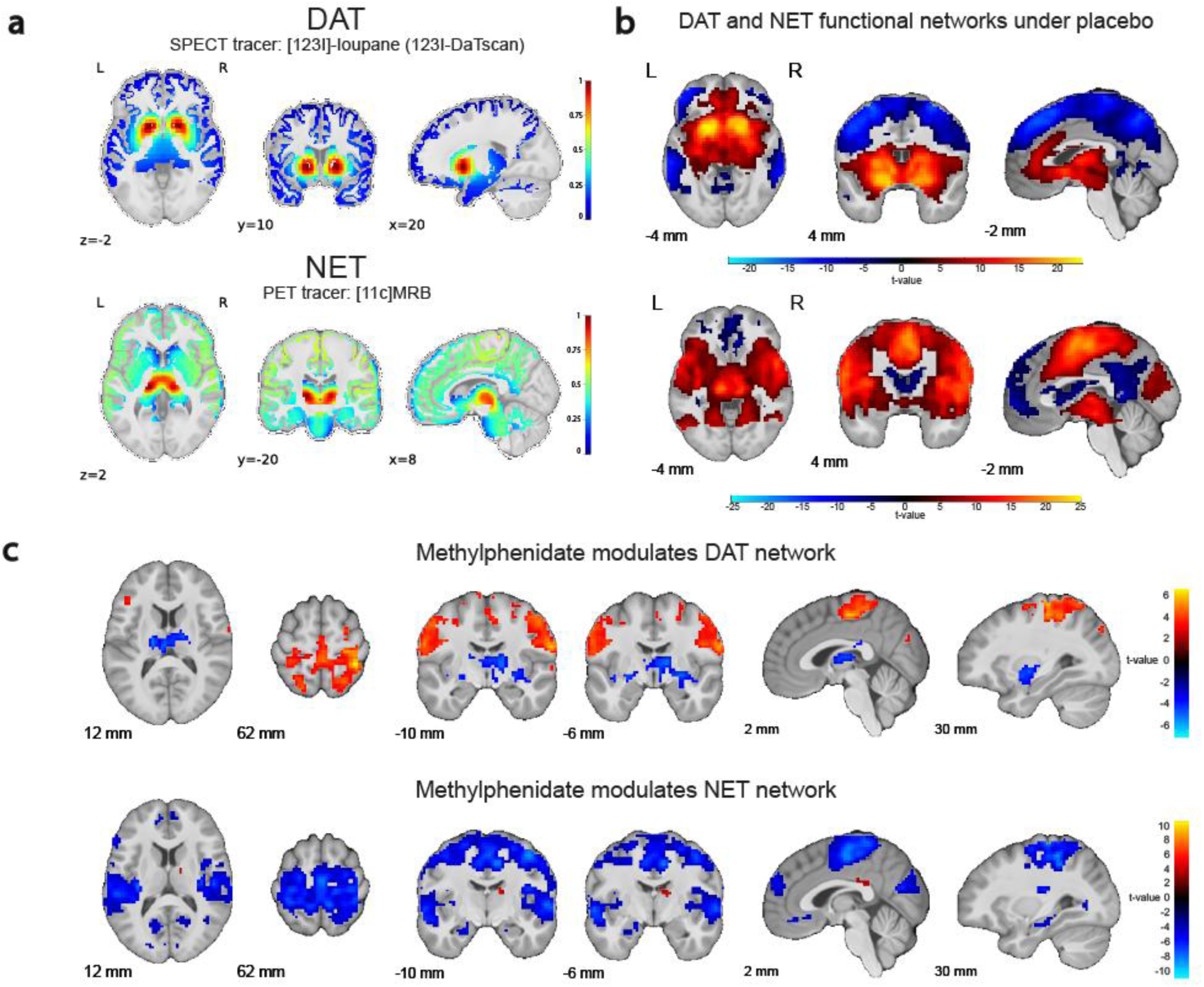
(a) Molecular templates of DAT (top) and NET (bottom) distribution. (b) DAT-enriched functional connectivity network (top) and NET-enriched functional connectivity network (bottom) under placebo. (c) Methylphenidate effect (versus placebo) on the functional connectivity networks related to DAT (top) and NET (bottom). For display purposes, the figures in panels b-c show data for voxels that passed the initial cluster forming threshold of p<0.001 without correction for multiple comparisons. The results are overlaid on the group-average T1-weighted anatomical scan in MNI152 coordinate space.

These analyses revealed dissociable functional connectivity maps for DAT and NET (Figure 1b). DAT-rich areas covaried positively with the striatum, vmPFC, and cingulate cortex, and negatively with large parts of the dorsal prefrontal and parietal cortex (placebo session; peak voxel positive: x y z = −17 10 −5, Z = Inf, p_FWE_clust_ = 1.941e-72, k = 8013 voxels; peak voxel negative: x y z = −43 −56 54, Z = Inf, p_FWE_clust_ = 2.776e-127, k = 18786 voxels). Conversely, NET-rich regions covaried positively with regions in the brain stem (centered on locus coeruleus) and large parts of the medial prefrontal cortex and insulae (peak voxel: x y z = 13 −20 1, Z = Inf, p_FWE_clust_ = 4.710e-146, k = 24332 voxels), and negatively with more focal regions in the anterior PFC, and posterior cingulate cortex (peak voxel biggest cluster, aPFC: x y z = 13 39 54, Z = Inf, p_FWE_clust_ = 5.480e-31, k = 2312 voxels).

Methylphenidate modulated both the DAT and NET-enriched functional connectivity networks (Figure 1c; Supplementary Table 1). Based on prior work (Dipasquale et al., 2020) this was anticipated for DAT but not for NET. Methylphenidate reduced DAT-enriched functional connectivity in subcortical areas, including the thalamus (peak voxel: x y z = −23 - 20 −5, Z = 4.90, p_FWE_clust_ = 2.697e-13, k = 500 voxels), while increasing DAT-enriched functional connectivity in dorsal cortical areas (peak voxel in superior parietal cortex: x y z = 39 −40 62, Z = 6.14, p_FWE_clust_ = 4.950e-32, k = 1737 voxels). By contrast, methylphenidate significantly decreased NET-enriched functional connectivity across widespread regions in cortex, with a peak effect in the insula (peak voxel in right insula: x y z = 39 −17 19, Z = Inf, p_FWE_clust_ = 2.109e-65, k = 5362 voxels).

Critically, as predicted, the effect of methylphenidate on the DAT-enriched functional connectivity network, but not the NET-enriched network, varied with interindividual differences in striatal dopamine synthesis capacity (Figure 2). Specifically, methylphenidate increased DAT-related connectivity in medial parietal cortex and in bilateral PFC clusters to a greater degree for participants with greater ventral striatal dopamine synthesis capacity (significant in parietal cortex, peak voxel: x y z = 6 −63 51, Z = 4.58, p_FWE_clust_ = 0.004, k = 61 voxels; left lateral PFC: x y z = −53 33 24, Z = 4.36; dorsal PFC: x y z = 29 26 57, Z = 4.40). The direction of this effect concurs with our (previously reported) observation that methylphenidate boosted neural RPE signaling in this same dataset also to a greater degree in people with higher dopamine synthesis capacity (van den Bosch et al., 2022). Remarkably, the effect of methylphenidate on DAT-enriched functional connectivity in the left lateral PFC (lPFC) correlated highly significantly with the effect of methylphenidate on RPE signaling in approximately the same region of lPFC (Figure 3; r = 0.288, p = 0.007; see van den Bosch et al. (2022)). The effect of methylphenidate on NET-enriched functional connectivity did not correlate with the drug effect on neural RPE signaling in the lPFC, neither in the cluster with a drug effect on DAT-related connectivity nor in the overlapping cluster with the previously reported drug effect on RPE signaling (r = 0.117, p = 0.286 and r = 0.060, p = 0.584, respectively).

**Figure 2.**
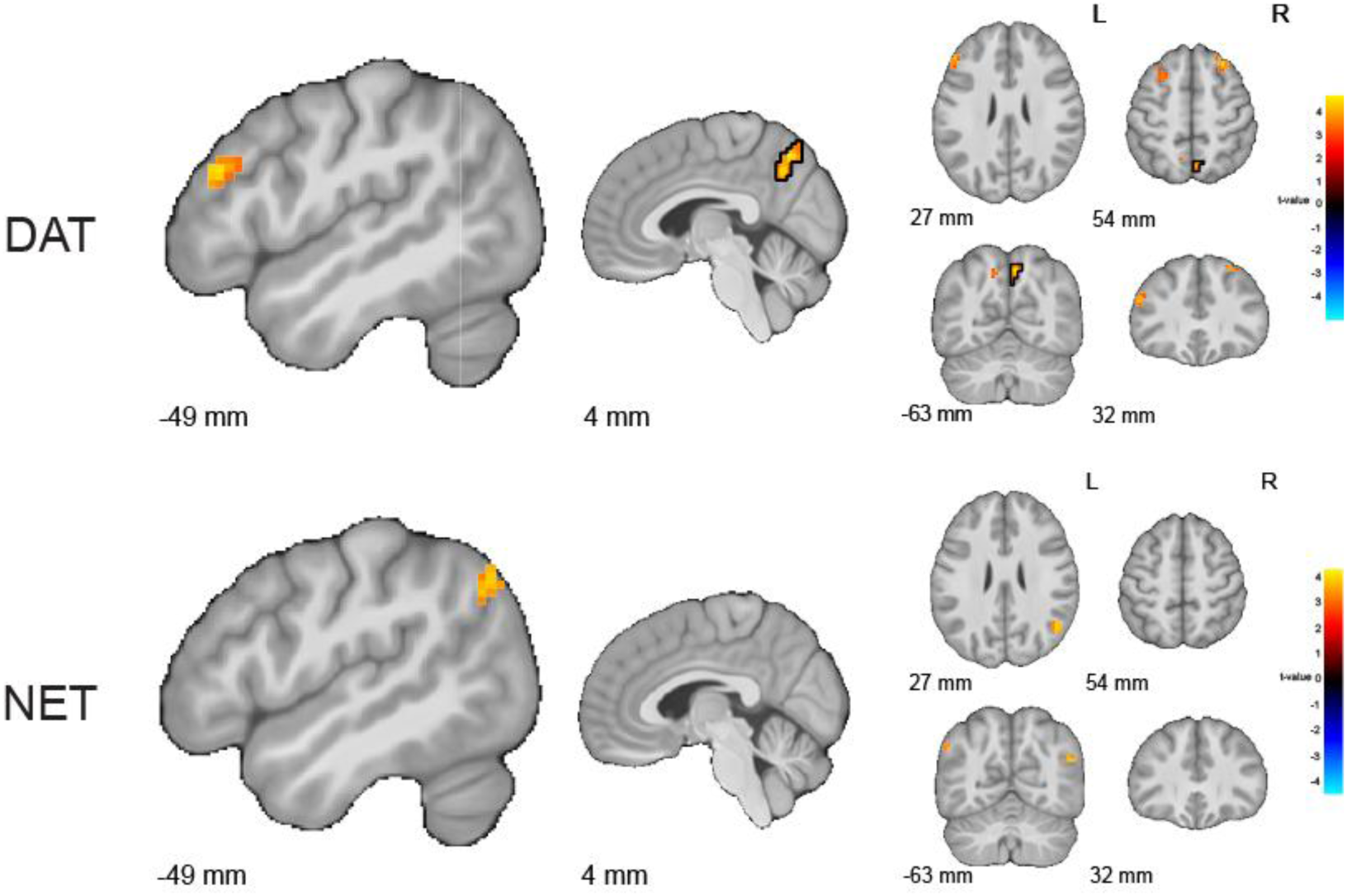
Methylphenidate effect on the functional connectivity networks related to DAT (top) and NET (bottom) as a function of individual differences in dopamine synthesis capacity in the ventral striatum. The effect of methylphenidate on DAT-enriched functional connectivity, but not NET-enriched functional connectivity, varied with ventral striatal dopamine synthesis capacity. Results are displayed with an uncorrected initial cluster-forming threshold of p<0.001. Clusters that are significant after family-wise error correction at p<0.05 are outlined with a black contour. The results are overlaid on the group-average T1-weighted anatomical scan in MNI152 coordinate space.

**Figure 3.**
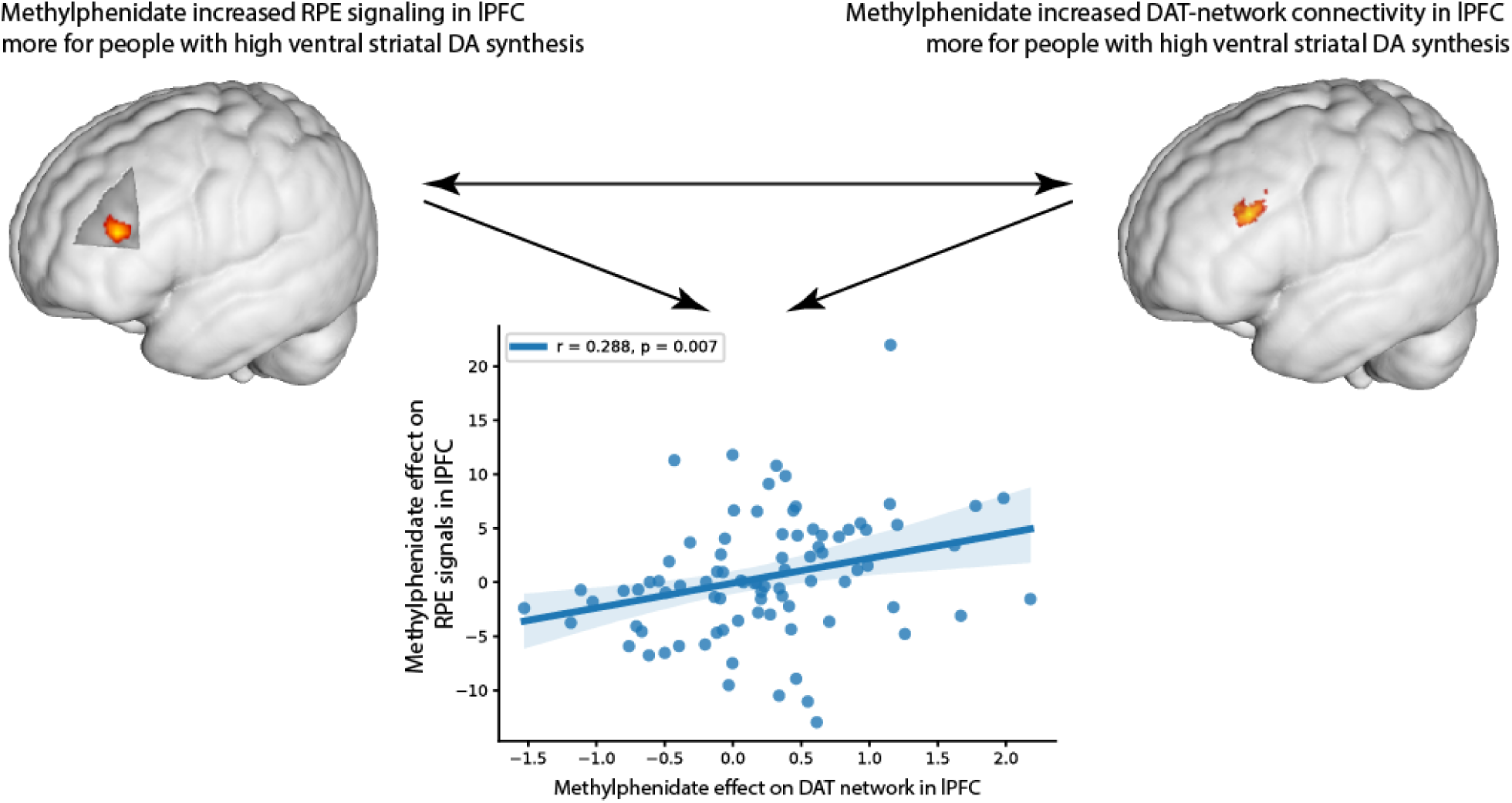
Striatal dopamine synthesis capacity correlates with methylphenidate effects on neural reward prediction error (RPE) signaling in a region of the prefrontal cortex (top left; van den Bosch et al. (2022)) that overlaps with the region that exhibits an association between striatal dopamine synthesis capacity and methylphenidate’s effect on DAT-enriched functional connectivity (top right). The scatterplot represents the correlation between the effects displayed in the top panels: The effect of methylphenidate on neural RPE signaling in the lPFC was positively correlated with the effect of methylphenidate on DAT-enriched functional connectivity in the lPFC.

Originally we had aimed to substantiate the specificity of our results to the specific drug targets by running the same target-enriched fMRI analyses using a spatial density map of SERT, derived from [^11^C]DASB PET imaging (Figure 4a). The SERT seemed a suitable control target, because methylphenidate binds only weakly to SERT compared to DAT and NET (Gatley et al., 1996; Luethi et al., 2018). The SERT-enriched functional connectivity map, however, overlapped greatly with the DAT-enriched connectivity map (Figure 4b; peak voxel: x y z = −10 −26 −5, Z = Inf, p_FWE_clust_ = 4.434e-84, k = 8023 voxels). Not surprisingly, the effect of methylphenidate on SERT-enriched functional connectivity was nearly identical to its effect on the DAT-enriched connectivity (Figure 4c). The drug decreased SERT-enriched functional connectivity in the thalamus (peak voxel: x y z = 6 −20 10, Z = 5.27, p_FWE_clust_ = 1.510e-14, k = 478 voxels), while increasing SERT-enriched connectivity in dorsal frontal and parietal cortex (peak voxel: x y z = 36 −40 62, Z = 5.67, p_FWE_clust_ = 3.808e-19, k = 652 voxels). Given the high collinearity between the DAT and SERT spatial similarity regressors (Figure 4d), we conclude that this prohibits further investigation of unique methylphenidate effects on DAT and SERT-enriched functional connectivity. The consequence of the spatial overlap between the DAT and SERT template maps is that REACT cannot be used to disentangle effects on DAT-enriched functional connectivity from those on SERT-enriched functional connectivity.

**Figure 4.**
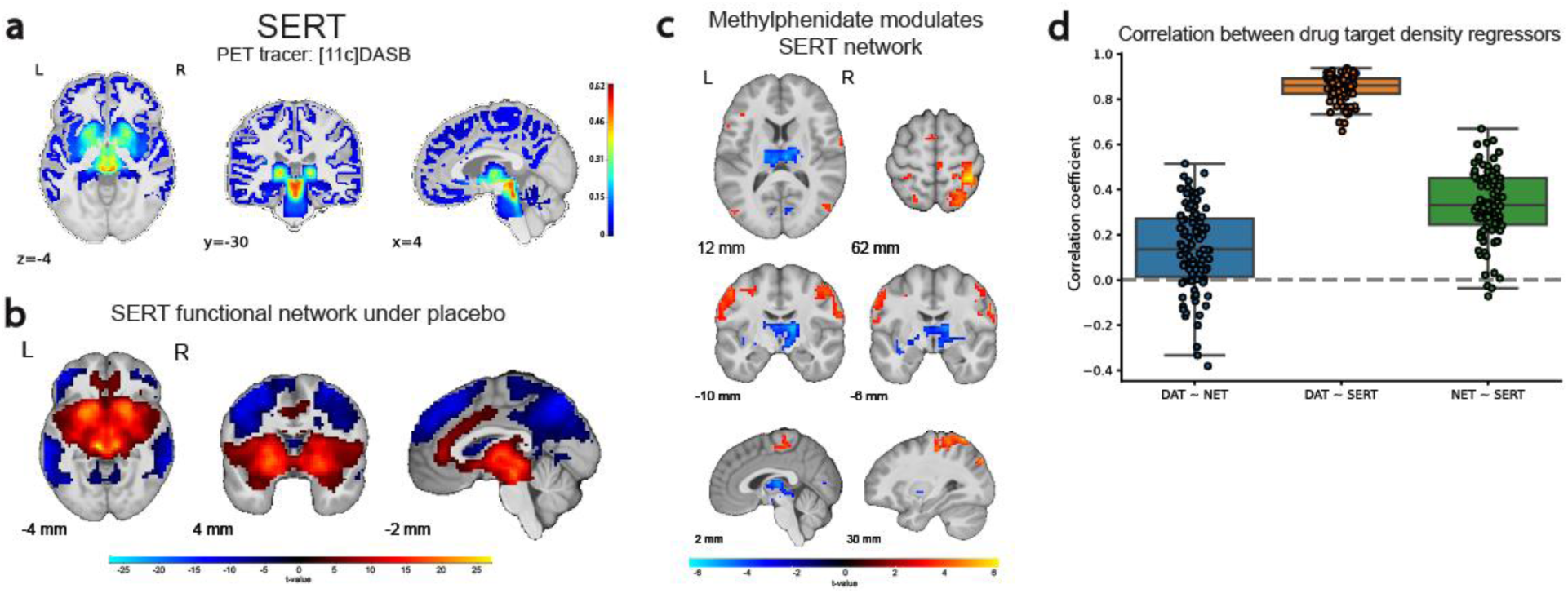
(a) Molecular template of SERT distribution. (b) SERT-enriched functional connectivity network under placebo. (c) Methylphenidate modulation of the SERT-enriched functional connectivity network. Results are displayed with an uncorrected initial cluster-forming threshold of p<0.001. The results are overlaid on the group-average T1-weighted anatomical scan in MNI152 coordinate space. (d) Pearson correlations between the spatial similarity regressors within participants.

## 4 Discussion

Methylphenidate is one of the most commonly used psychoactive drugs, but the molecular mechanisms by which it elicits its cognitive effects have not been elucidated. One challenge is that methylphenidate can affect specific cognitive functions and associated brain signals by modulating the dopamine system, via its action on DAT, and/or the noradrenaline system, via action on NET. A recently developed dual-regression approach (REACT; Dipasquale et al. (2019)) promises to hone in on the molecular mechanisms underlying changes in BOLD signal by enriching fMRI analyses with information about the spatial distribution of molecular targets of interest. In one of the first applications of the method methylphenidate was observed to affect DAT-enriched but not NET-enriched functional connectivity, measured during resting state (Dipasquale et al., 2020). Reliance on REACT for inferring molecular target specificity is increasingly common, in part because of the availability of large resting state fMRI datasets and the ease of linking these with (publicly available) PET templates of molecular targets. However, despite the promise in principle of this approach, the molecular specificity of the method, and thus its suitability for disentangling the role of dopamine from noradrenaline signaling, has not yet been firmly established.

We leveraged a unique opportunity to assess the molecular and functional validity of REACT for disentangling the role of dopamine from that of noradrenaline by using an existing pharmaco-fMRI dataset from a large sample of healthy volunteers. In a previous analysis of this dataset we had already established methylphenidate effects on a canonical functional signature of dopamine: neural RPE signaling. Moreover, the importance of dopamine for this effect of methylphenidate on RPE signaling was substantiated by the finding that it varied with interindividual variation in striatal dopamine synthesis capacity (van den Bosch et al., 2022). In the current study, we reanalyzed those data using REACT and found clear and distinct effects of methylphenidate on DAT-enriched and NET-enriched functional connectivity networks. Importantly, only the drug effect on DAT-related connectivity, but not NET-related connectivity, varied as a function of striatal dopamine synthesis capacity. Furthermore, the methylphenidate effect on DAT-enriched connectivity in the lPFC was significantly associated with the drug effect on neural RPE signaling in the lPFC, and both of those drug effects varied with differences in dopamine synthesis capacity in the ventral striatum. These results provide functional validation of the method, because they dissociate methylphenidate effects on DAT from NET not simply during resting state but also as a function of neural RPE signaling. In addition, they demonstrate that the method confers molecular specificity by revealing that only the effects on DAT-enriched functional connectivity were associated with dopamine synthesis capacity, as measured with [^18^F]FDOPA PET.

The finding that methylphenidate’s effects on neural RPE signaling in PFC are accompanied by effects on DAT-enriched functional connectivity in the same location substantiates the conclusion that methylphenidate enhances the expression of frontal function by boosting the striato-frontal transmission of information outflow from basal ganglia-centered DAT-rich regions to PFC. The observation that, across all participants, methylphenidate decreased functional connectivity between DAT-rich regions, such as the basal ganglia and the thalamus, while increasing it with cortex is tantalizingly reminiscent of pervasive functional models of cortico-striato-thalamic pathways (Alexander & Crutcher, 1990; Mink, 1996), according to which increases in striatal dopamine disinhibit the cortex by suppressing the constant inhibitory influence from the output nuclei of the basal ganglia on the thalamus. The degree to which the drug achieved its impact on frontal function, however, depended on baseline levels of dopamine synthesis capacity, perhaps due to greater dynamic range for effects of transporter blockade in high-dopamine participants.

In addition to validating the tool for disentangling the role of DAT and NET in psychostimulant effects on brain function, we demonstrate that the method is not suitable for dissociating effects on DAT-enriched connectivity from those on SERT-enriched connectivity, highlighting a critical limitation of the approach. This limitation is grounded in the fact that the method leverages the correspondence of spatial distributions of molecular targets with spatial patterns of whole-brain BOLD signal at any given timepoint, measured with MRI. Therefore, REACT can only discriminate those targets for which distributional differences are discernable at the limited spatial resolution of MRI. Thus, even if the spatial distributions of DAT and SERT are discriminable using techniques with higher spatial resolution (e.g. transcriptomics or postmortem autoradiography (Amunts et al., 2020; Fazio et al., 2016)), the spatial resolution of MRI is not sufficient to resolve those differences. Our conclusion that the effects of methylphenidate on DAT-related functional connectivity reflect drug action on the DAT and not the SERT is based on prior evidence that methylphenidate has relatively little effect on the serotonin system compared with the dopamine system (Faraone, 2018) and the fact that methylphenidate has much higher affinity for DAT than SERT (Gatley et al., 1996; Pan et al., 1994).

Unlike the DAT and SERT functional connectivity networks, the DAT and NET maps were not similar, which enabled dissociating effects of methylphenidate on the DAT network from those on the NET network. The discriminability of the DAT and NET functional networks itself does not constitute evidence for their molecular specificity, but did encourage us to assess their respective associations with dopamine synthesis capacity, measured with PET in the same individuals. One implication of the unique association between the drug effect on DAT-related connectivity and individual-level [^18^F]FDOPA uptake is that the method might be used for isolating the unique role of dopamine in catecholaminergic drug effects using simple pharmaco-fMRI designs. That would complement more complex comparative multi-drug designs (e.g. to compare effects of methylphenidate with atomoxetine) that have hitherto been necessary for making inference about dopamine-specificity (Chen et al., 2024; Cremer et al., 2023; Schulz et al., 2017).

The methylphenidate effects on DAT/NET-enriched functional connectivity that we observed were more widespread than those observed in a previous study (Dipasquale et al., 2020), in which methylphenidate affected DAT-related functional connectivity only in small clusters in the pre- and postcentral gyri and supramarginal gyrus, and no effects on NET-related functional connectivity were observed. We observed greater changes in DAT-enriched functional connectivity, but in similar cortical locations, with additional subcortical effects. In addition, we observed extensive effects of methylphenidate on the NET-related functional connectivity. The drug decreased NET-related connectivity throughout the frontal and parietal parts of the network. This reduction in large-scale network coherence might reflect a NET-mediated drug effect on the sustained attention required to keep performing the task (Berridge et al., 2012; Marshall et al., 2019). Thus, one possibility is that these differences between studies reflect the fact that the current fMRI data were acquired during task performance, while Dipasquale et al. (2020) reported on networks measured during resting state. Different brain networks are recruited depending on task demands versus rest and effects of chemical neuromodulators are well established to depend on the baseline activity state of the target networks. Hence, administration of psychoactive drugs would also have influenced those networks differently. Such differences were also observed in a study investigating the effect of atomoxetine on functional connectivity as measured with magneto-encephalography (Pfeffer et al., 2021). Cortical circuit modeling was used to demonstrate that even subtle differences in the baseline excitatory/inhibitory state of the system can result in qualitatively different drug effects on functional connectivity. Thus, methylphenidate might well have elicited larger-scale decreases in NET-related functional connectivity in the current versus the previous study, because increases in noradrenaline boost the neural gain and cortical efficiency of the attention network to a greater degree when the baseline state is more active. That would be consistent with the sigmoid shaped relationship between signal detection performance and catecholamines in classic network models of catecholamine function (Servan-Schreiber et al., 1990). Having said this, the current study was not set up or optimized for isolating the role of noradrenaline in methylphenidate’s effects. Future work is necessary, for example using independent individual-level NET density indices of noradrenaline signaling from PET imaging, to substantiate the hypothesis that the drug effects on the large scale NET network indeed reflect selectively noradrenaline signaling.

## 5 Data and Code Availability

The minimally processed data used in this study and the overarching project it is part of are available from the Radboud Data Repository (https://doi.org/10.34973/wn51-ej53; custom data use agreement RU-DI-HD-1.0). The final data derivatives relevant to the current work, as well as all code for data analysis, are available from a separate collection on the Radboud Data Repository (https://doi.org/10.34973/zve0-ym39).

## 6 Author Contributions

Conceptualization: R.C., R.B.; analysis: R.B.; software: R.B.; writing: R.B.; review: R.B., R.C.

## Supplementary information

**Supplementary Table 1.**
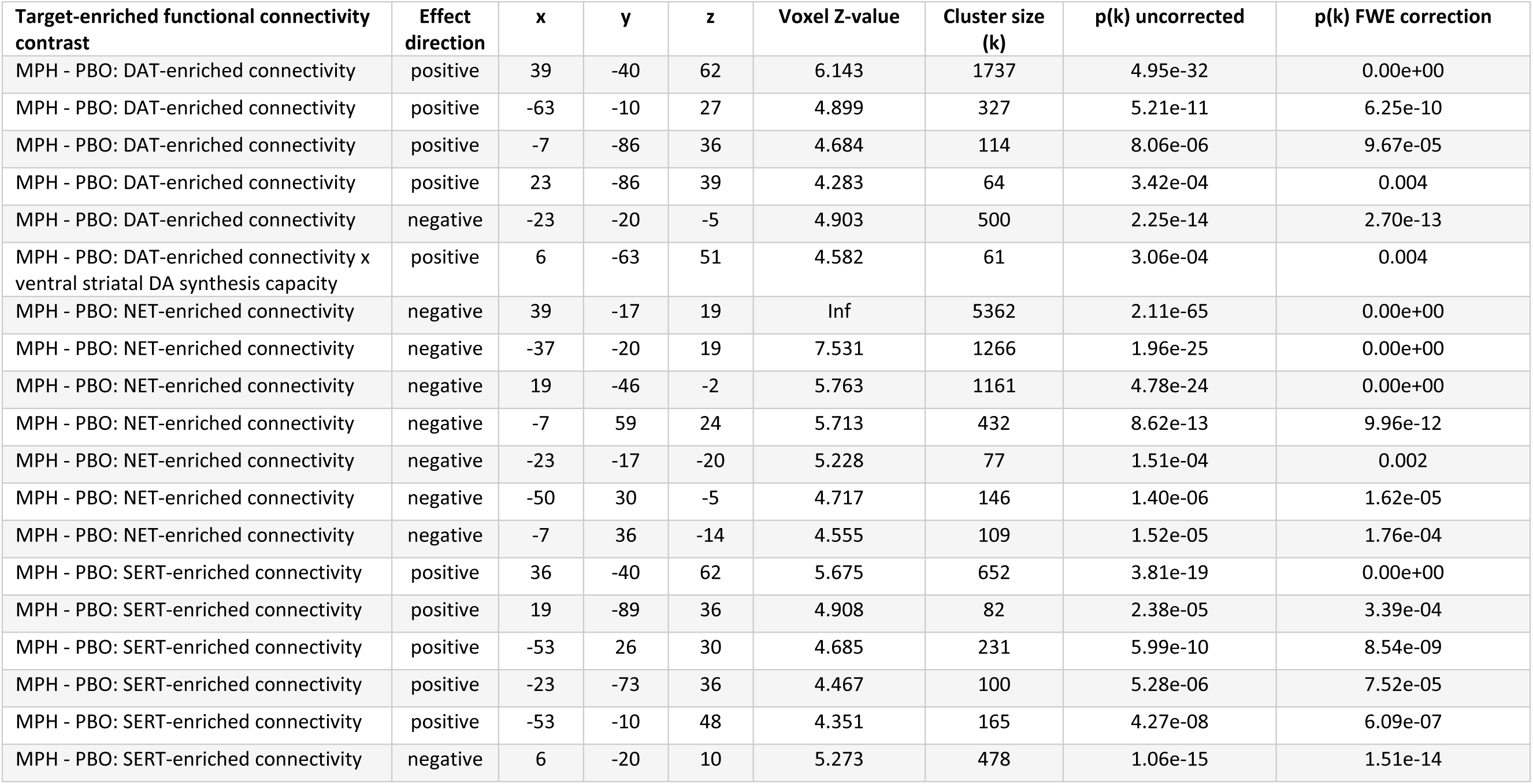
List of all clusters with effects of methylphenidate at p < 0.001 uncorrected and/or p < 0.05 with cluster-level FWE correction at the whole-brain level. The x y z coordinates are in MNI152 coordinate space. MPH: methylphenidate; PBO: placebo; DAT: dopamine transporter; NET: noradrenaline transporter; SERT: serotonin transporter.

