## Supplementary Table 1 for "Validating molecular target-enriched fMRI for disentangling drug effects on dopamine"

### Supplementary information

Supplementary Table 1. List of all clusters with effects of methylphenidate at $p < 0.001$ uncorrected and/or $p < 0.05$ with cluster-level FWE correction at the whole-brain level. The x y z coordinates are in MNI152 coordinate space. MPH: methylphenidate; PBO: placebo; DAT: dopamine transporter; NET: noradrenaline transporter; SERT: serotonin transporter.

| Target-enriched functional connectivity contrast | Effect direction | x | y | z | Voxel Z-value | Cluster size (k) | p(k) uncorrected | p(k) FWE correction |
|---|---|---|---|---|---|---|---|---|
| MPH - PBO: DAT-enriched connectivity | positive | 39 | -40 | 62 | 6.143 | 1737 | 4.95e-32 | 0.00e+00 |
| MPH - PBO: DAT-enriched connectivity | positive | -63 | -10 | 27 | 4.899 | 327 | 5.21e-11 | 6.25e-10 |
| MPH - PBO: DAT-enriched connectivity | positive | -7 | -86 | 36 | 4.684 | 114 | 8.06e-06 | 9.67e-05 |
| MPH - PBO: DAT-enriched connectivity | positive | 23 | -86 | 39 | 4.283 | 64 | 3.42e-04 | 0.004 |
| MPH - PBO: DAT-enriched connectivity | negative | -23 | -20 | -5 | 4.903 | 500 | 2.25e-14 | 2.70e-13 |
| MPH - PBO: DAT-enriched connectivity x ventral striatal DA synthesis capacity | positive | 6 | -63 | 51 | 4.582 | 61 | 3.06e-04 | 0.004 |
| MPH - PBO: NET-enriched connectivity | negative | 39 | -17 | 19 | Inf | 5362 | 2.11e-65 | 0.00e+00 |
| MPH - PBO: NET-enriched connectivity | negative | -37 | -20 | 19 | 7.531 | 1266 | 1.96e-25 | 0.00e+00 |
| MPH - PBO: NET-enriched connectivity | negative | 19 | -46 | -2 | 5.763 | 1161 | 4.78e-24 | 0.00e+00 |
| MPH - PBO: NET-enriched connectivity | negative | -7 | 59 | 24 | 5.713 | 432 | 8.62e-13 | 9.96e-12 |
| MPH - PBO: NET-enriched connectivity | negative | -23 | -17 | -20 | 5.228 | 77 | 1.51e-04 | 0.002 |
| MPH - PBO: NET-enriched connectivity | negative | -50 | 30 | -5 | 4.717 | 146 | 1.40e-06 | 1.62e-05 |
| MPH - PBO: NET-enriched connectivity | negative | -7 | 36 | -14 | 4.555 | 109 | 1.52e-05 | 1.76e-04 |
| MPH - PBO: SERT-enriched connectivity | positive | 36 | -40 | 62 | 5.675 | 652 | 3.81e-19 | 0.00e+00 |
| MPH - PBO: SERT-enriched connectivity | positive | 19 | -89 | 36 | 4.908 | 82 | 2.38e-05 | 3.39e-04 |
| MPH - PBO: SERT-enriched connectivity | positive | -53 | 26 | 30 | 4.685 | 231 | 5.99e-10 | 8.54e-09 |
| MPH - PBO: SERT-enriched connectivity | positive | -23 | -73 | 36 | 4.467 | 100 | 5.28e-06 | 7.52e-05 |
| MPH - PBO: SERT-enriched connectivity | positive | -53 | -10 | 48 | 4.351 | 165 | 4.27e-08 | 6.09e-07 |
| MPH - PBO: SERT-enriched connectivity | negative | 6 | -20 | 10 | 5.273 | 478 | 1.06e-15 | 1.51e-14 |
